# Developing a model to implement marker-assisted selection for root-knot nematode resistance in common bean

**DOI:** 10.1101/2025.01.31.635933

**Authors:** Talissa de Oliveira Floriani, Henrique Castro Gama, Bruna Marques Moreno, Guilherme Alexandre Luz Costa, Willian Giordani, Alisson Fernando Chiorato, Líllian Bibiano, Travis A. Parker, Luis Eduardo Aranha Camargo, Maria Lucia Carneiro Vieira, Antonio Augusto Franco Garcia

**Author notes:** Corresponding author: Departamento de Genética, Escola Superior de Agricultura Luiz de Queiroz, Universidade de São Paulo, 13418-900 Piracicaba, Brazil. Authors who contributed equally to the study (Talissa de Oliveira Floriani, Henrique Castro Gama).

## Abstract

Common bean (*Phaseolus vulgaris* L.) is a vital crop for direct human consumption, with essential nutrients and valuable protein that provides food security in developing countries. However, its cultivation faces significant threats from *Meloidogyne incognita*, a root-knot nematode (RKN), resulting in considerable yield loss. Developing crop resistance remains a key strategy for mitigating nematode infections. To investigate the genetic architecture of common bean responses to RKN (specifically, race 3 of *M. incognita*), we performed controlled crosses between the genotypes ‘IAC-Tybatã’ and ‘Branquinho’ with contrasting resistance. The resulting segregating population (*F_2_*) of 333 individuals was genotyped using GBS (genotyping-by-sequencing). We used a phenotyping approach, already optimized in the lab, to collect trait data for a subset of 200 F_2:3_ families. Evaluations of egg mass (EM), root-galling index (GI), and root dry mass (RM) were conducted 30 days after RKN inoculation under greenhouse conditions, in a completely randomized design with ten replicates. Linkage and quantitative trait loci (QTL) mapping were performed, while functional mapping of associated regions facilitated identification of candidate genes. A linkage map encompassing 954 SNPs assigned to 11 linkage groups totaling 1,687 cM formed the basis for Interval Mapping (IM), Composite Interval Mapping (CIM), and Multiple Interval Mapping (MIM), revealing four major QTLs (on Pv03, Pv05, Pv08, and Pv10) and epistasis between QTL on Pv08 and on Pv10 associated with the GI trait. No significant QTL were identified for EM and RM. The model enabled calculation of genotypic values through marker-assisted selection (MAS). The high correlation between observed and predicted values (0.72) underscores the model’s significance. Candidate genes previously associated with nematode resistance were also identified within the QTL interval on chromosome Pv10. Our results will be valuable for future selection of varieties resistant to this important crop disease.

## Introduction

The common bean (*Phaseolus vulgaris* L., Fabaceae) is the most important grain legume for direct human consumption globally, a source of dietary proteins, fibers, and nutrients, especially in developing countries (Akibode and Maredia 2012; Bitocchi et al. 2017; Joshi and Rao 2017). With over 800 million people facing hunger, sustainable and high yield agricultural practices are urgently needed. In 2022, the global production of primary crop commodities, including common beans, reached approximately 9.6 billion tons, an increase of 0.7% since 2021. Brazil, the third-largest common bean producer, provides over 2.8 million tons (FAO 2023). Originating in central Mexico and the southern Andes, the crop was domesticated differentially, yielding two distinct gene pools: the Andean and Mesoamerican, the latter containing greater genetic diversity (Gepts et al. 1986; Gepts and Papa 2003; Kwak and Gepts 2009; Bitocchi et al. 2012, Schmutz et al. 2014). In Brazil, the most consumed varieties are medium to small-sized black and carioca beans (Blair et al. 2013). This genetic diversity is vital for food security and agricultural sustainability.

Common bean cultivation faces severe phytosanitary challenges, including those arising from nematode attacks, which compromise the root system and impair plant growth, and leave the crop more susceptible to biotic and abiotic stresses (Grundler and Hofmann 2011). Cyst nematodes (*Heterodera* spp.), lesion nematodes (*Pratylenchus* spp.), and root-knot nematodes (RKN; *Meloidogyne* spp.) are collectively significant as primary pathogenic genera, drastically impacting production, with annual losses of up to US$ 173 billion (Gamalero and Glick 2020). Of particular concern are RKN, endoparasites that establish their feeding sites intricately, inducing differentiation in host root cells, which ultimately leads to the formation of galls (Jones and Goto 2011). In common bean cultivation, the most prevalent and destructive RKN species, *M. incognita* and *M. javanica*, are responsible for the development of galls (Singh and Schwartz 2011; Santos et al. 2012). Galls emerge as a consequence of abnormal growth triggered by gall-inducing organisms. The resultant root damage, induced by *M. incognita*, adversely impacts water and nutrient absorption in infected plants, resulting in symptoms such as dwarfism and suboptimal development of the vegetative system, which ultimately compromises the overall yield (Santos et al. 2012). The global impact of RKN induced reductions in productivity is considerable, with estimated losses of around 10% (Whitehead 1998).

Especially in tropical and subtropical regions, the significant damage caused by the nematode is attributed to its ability to suppress the host defense, leading to high population densities (Sikora et al. 2018). Chemical control methods, including fumigant nematicides, have been used in a number of countries and remain the main strategy for managing plant-parasitic nematodes (Desaeger et al. 2020). Although nematicides are an alternative RKN control method, their use has provoked environmental concerns. The most effective approach to prevent RKN-related diseases is cultivating resistant cultivars (Barbary et al. 2015). Systematically evaluating available germplasm to identify genetic resistance sources is a crucial breeding program strategy, to minimize the damage caused by RKNs. Despite associated challenges (for example, the difficulty of measuring phenotypes for a root characteristic related trait), notable findings have been published on the response of common bean genotypes to RKN and the discovery of putative resistance genes (Ferreira et al. 2010; Costa et al. 2019; Giordani et al. 2022; Dias et al. 2023).

The advent of SNP-based markers and availability of annotated genome sequences have accelerated the association of quantitative trait loci (QTL) and candidate genes. Mapping QTL resistance to RKN is reported in various species, including soybean (*Glycine max*), cowpea (*Vigna unguiculata*) (Shearin et al. 2009; Huynh et al. 2016; Leal-Bertioli et al. 2016; Santos et al. 2018; Ndeve et al. 2019), sweet potato (*Ipomoea batatas*) (Oloka et al. 2021), and carrot (*Daucus carota*) (Parsons et al. 2015). In terms of nematode resistance in the common bean, data is scarce, with most studies focused on *Heterodera glycines* (Shi et al. 2021). To our knowledge, there is only one study of RKN resistance using a common bean diversity panel (Giordani et al. 2022), wherein genotype-phenotype associations were clearly identified. The SNP effects were subsequently validated through traditional QTL mapping in a BC_2_F_4_ population, revealing a major QTL on Pv05, eight lesser genomic regions on different chromosomes, and 216 candidate genes associated with the host response. These included 14 resistance gene analogs (RGA), and five genes detected with a differential expression in a previous transcriptome analysis (Santini et al. 2016).

Traits associated with RKN disease pose significant control challenges, as they are time-consuming and expensive to evaluate and breed. Marker-assisted selection (MAS) is an opportunity for breeders to streamline the process of creating resistant cultivars. While the comparative results between genomic selection (GS) and MAS are still debated, both approaches remain ideally suited to different purposes, especially in relation to complex traits. GS is considered one of the best methods for predictive breeding, particularly for traits controlled by small-effect QTL (Bernardo and Yu 2007). Conversely, MAS is highly effective for disease resistance, focused on a limited number of QTLs and traits. For example, MAS has proven one of the best approaches to improving resistance to leaf rust in wheat, with high predictive accuracy (Beukert et al. 2020). Successful applications of MAS include soybean cyst nematode resistance, in which identification and transfer of the resistance gene *Rhg1* were effectively achieved (Cregan et al. 1999; Young 1999; Marcelino-Guimarães et al. 2007). Importantly, Chaiprom et al. (2024) suggest additional genes that may enhance *Rhg1*-mediated SCN resistance. Markers associated with resistance to common bean bacterial blight and anthracnose (Garzón et al. 2008) have shown the utility of MAS (Yu et al. 2000; O’Boyle et al. 2007).

The primary objective of this study was to advance our understanding of the genetic architecture underlying traits associated with the common bean response to RKN and develop a strategy to efficiently implement MAS. Part of our focus was to map QTL, to estimate their location and effects. We also investigated an MAS model which facilitates an estimate of the genetic values of individuals based on the identified QTL. Additionally, to advance knowledge of the trait, we researched candidate genes related to RKN resistance. This work marks a significant advance in unraveling the genetic foundations of RKN resistance in common beans, laying the groundwork for implementing MAS in breeding populations to enhance pathogen resistance in the bean family.

## Material and methods

### Plant material

An *F*_2_progeny was used, consisting of 333 plants derived from a single cross between the inbred lines ‘IAC-Tybatã’ (moderately resistant) and ’Branquinho’ (susceptible) (TB population), both accessions of Mesoamerican origin. They belong to the common bean germplasm bank of The Agronomic Institute (IAC), Campinas, Brazil, where the crosses were developed. In addition to the contrast in terms of their response to RKN infection (Giordani et al. 2022), the lines differ in angular leaf spot and anthracnose response, plant architecture, and phenological cycle (Perseguini et al. 2011; Diniz et al. 2018). Furthermore, both accessions belong to the ’carioca’ bean type, primarily cultivated in Brazil, representing around 70% of Brazilian consumption (Souza et al. 2020). Branquinho is a landrace, known for its tolerance to slow darkening in grain storage silos (Chiorato et al. 2020), a trait of high commercial value (Elsadr et al. 2011; Rodrigues et al. 2019). IAC-Tybatã, meanwhile, is a high-yielding elite cultivar developed through the IAC breeding program 24 years ago, with resistance to viruses caused by bean golden mosaic virus (BGMV) and bean common mosaic virus (BCMV), anthracnose, and rust caused by *Colletotrichum lindemuthianum* and *Uromyces appendiculatus*, respectively. The *F*_2_ progeny was used to construct the linkage map. An *F*_2:3_ segregating population of 200 lines, produced by selfing *the F*_1_ ∧ *F*_2_ plants, was used to phenotype the response to RKN inoculation and QTL mapping.

### Root-knot nematode resistance phenotyping

The average phenotypic value of the *F*_2:3_ progeny was used to estimate the phenotypic value of *F*_2_ plants, increasing the precision of quantitative trait loci detection (Zhang and Xu 2004). The parental lines, the *F*_2:3_ families, and the control genotypes ’Puebla- 152-CIAT’ (moderately resistant) and ’Jamapa’ (susceptible) were evaluated for reaction to *M. incognita* race 3 infection using the Atamian et al. (2012) protocol, with minor modifications, as described in Giordani et al. (2022). Phenotypic evaluations were carried out using a completely randomized experimental design with ten repetitions.

Seeds were pre-germinated in an incubator at 26 °C until the roots reached 1 to 2 cm in length and were transferred to plastic zipper bags (24 cm × 17 cm) containing germination paper. The bags were put into plastic boxes, kept in a greenhouse and watered daily with distilled water.

The inoculum was raised by inoculating the susceptible tomato line ’Santa Clara VF5600’ with *M. incognita* race 3 and harvested 60 days after inoculation. Nematode eggs were extracted from the roots, which were washed, cut, and manually stirred for 2 min in 500 mL of 12% NaCl solution in a bottle. The solution was sieved through 425-, 90-, and 25 µm meshes to retain the eggs, and washed with distilled water to remove the excess of sodium chloride.

Subsequently, they were deposed on double-layer disposable wipes fitted over a metal basket inside a Petri dish and incubated for 5 to 8 days. Freshly hatched juvenile nematodes (J2) (infective phase), passed through the paper, were collected and counted using a Peters slide under an optical microscope. An inoculation solution was prepared by dilution, resulting in a final concentration of 300 J2/mL. The plants were placed horizontally before inoculation and the roots inoculated with approximately 1,500 J2s (*M. incognita* juveniles - Stage 2) in 5 mL of the suspension and kept in a greenhouse, irrigated daily with Hoagland’s nutritive solution (Hoagland and Arnon 1950). Thirty days after inoculation, the number of egg masses (EM), root-galling index (GI), and dry root mass (RM) were estimated, to assess the response to RKN infection. Egg masses were dyed and counted under a stereoscope (10x magnifier), after infusing the roots with 15 mL of eriglaucine (75 mg/L) for 12 h (Suplementary Figure 5). The Bridge and Page (1980) scale was used to index the root-galling symptoms, and the dry root mass was recorded after drying the roots in a conventional drying kiln at 90 °C for four hours, weighed on an analytical balance.

### Phenotypic data analysis

The statistical model to analyze RKN resistance phenotypic data was:

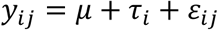

in which, *y*_j*k*_ is the observed phenotypic value of the *i* genotype in repetition *j*; µ is the intercept; *τ*_𝑖_ is the random effect of genotype *i*, with *τ*_𝑖_𝑁(0, *σ*_τ_^2^); and 𝜀_𝑖j_ is the experimental error associated with the *i* genotype in repetition *j*, 𝜀_𝑖j_*N*(0, *σ*^2^).

The best linear unbiased predictors obtained for trait effects were used to estimate Pearson’s correlation coefficient between traits. Given the unbalanced nature of the data, due to the natural difficulty of phenotyping for nematode reactions, the heritability was calculated following Cullis et al. (2006):

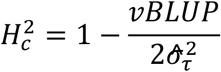

where, *H*_c_^2^ is heritability; 𝑣𝐵𝐿𝑈𝑃 the variance of the average difference between the two best linear unbiased predictors, and *σ*^_τ_^2^ the genetic variance. The analysis was carried out using ASReml-R (Butler et al. 2009) in the R statistical environment.

### Genotypic data

Both parents and 333 *F*_2_plants were genotyped using the GBS protocol (Poland et al. 2012). For genomic library assembling, the DNA was extracted from young leaves using the CTAB protocol at 2% (Doyle and Doyle 1990), purified, normalized to 30 µg/mL, and digested with *Pst*I and *Mse*I enzymes. Subsequently, barcode adapters were ligated to the fragments, combined, and amplified by PCR (Elshire et al. 2011). The library was purified and resuspended. BioAnalyzer DNA analysis was performed, to precisely measure the size and concentration of DNA fragments and smears. KAPA Library Quantification Kits were used for accurate, reliable, and reproducible qPCR-based library quantification.

Sequencing was run on the Illumina NextSeq 2000 platform, with Illumina NextSeq 1000/2000 P2 Reagents (100 cycles) v3, at the Functional Genomic Center at the Luiz de Queiroz College of Agriculture, Piracicaba, Brazil. GBS-SNP calling was performed using the default parameters of the software package TASSEL version 5.0 (Bradbury et al. 2007). The tags were aligned with the reference genome of *Phaseolus vulgaris* v. 2.1 (Schmutz et al. 2014) available in the Phytozome (https://phytozome-next.jgi.doe.gov/) platform using Bowtie 2 (Langmead and Salzberg 2012). Heterozygous SNPs with missing data > 20%, depth < 10%, and minor allele frequency (MAF) < 0.2 were removed using VCF tools (Danecek et al. 2011).

### Genetic map construction

The genotypic data of the *F*_2_progeny (comprising 2169 markers) were tested for the expected Mendelian frequencies (1:2:1), and those exhibiting segregating distortions were excluded after correcting for multiple tests. The recombination fraction was initially estimated by two-point analysis to form linkage groups, considering a maximum recombination fraction of 0.5 and a LOD Score of 6.573, based on the number of markers and tests. Distances between markers were used to order markers within each linkage group, guided by the reference genome. The Kosambi function was used to transform the recombination fractions into distances in cM (Kosambi 1943). The ordering distances were estimated based on the multipoint approach of a hidden Markov model. The marker order was inspected and checked based on heatmap graphs. The genetic linkage map construction was performed using OneMap v.2.1.3 (Margarido et al. 2007).

### QTL mapping and marker-assisted selection

QTL analysis was carried out regarding the egg mass (EM), root-galling index (GI), and root dry mass (RM). A linkage map was used to estimate each QTL genotype multipoint probability at step sizes of 1 cM. A single QTL model from interval mapping (IM) was applied as an initial approach, to estimate the QTL number and positions using the EM algorithm (Expectation-Maximization) and the Haley-Knott regression to adjust the model (Haley and Knott 1992). The significance of QTL effects was tested based on 5000 permutations (Churchill and Deorge 1994). Furthermore, a CIM model (Composite Interval Mapping) (Zeng 1994) was used to refine the results after selecting markers as covariates using a multiple regression approach, considering a window size of 15 cM. The CIM model was:

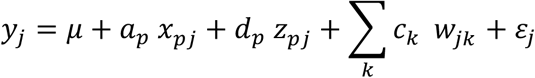

In which: *y*_j_is the phenotypic mean of individual 𝑗; 𝜇 is the intercept; 𝑎_𝑝_ is the additive of the QTL in the interval tested; 𝑑_𝑝_ is the dominance effect of the QTL in the interval tested, 𝑥_𝑝j_ indicator variable for the additive effect of the QTL for individual 𝑗; 𝑧_𝑝j_ in-dicator variable for the dominance effect of QTL for individual 𝑗; ∑_*k*_ 𝑐_*k*_ 𝑤_j*k*_ cofactor terms representing background markers outside the target interval to control for genetic background effects. The indicator variables𝑥_𝑝j_ and 𝑥_𝑞j_ are coded based on the genotype of the QTL for each individual 𝑗, with values of 1 for genotypes with QTL QQ, 0 for QTL genotypes of Qq or -1 if the QTL is qq, respectively; and 𝜀_j_is the residual of the individual where 𝜀_j_∼*N*(0, *σ*^2^).

The results of the CIM model were used to develop a more comprehensive mapping framework by multiple interval mapping (MIM) (Kao et al. 1999, 2002; Zeng et al. 1999) with 1000 permutation tests conducted to define the significance threshold of QTL detection. The search strategy for the final model was established with the step- wise function, to perform the stepwise regression; within the estimation procedure, the model was tested multiple times, adding each QTL per time point and comparing the P values summary from each model using the functions *makeqtl*, *fitqtl*, and *refineqtl.* After no more QTL were added, we tested for the presence of epistatic interactions between QTLs. The selection criteria for selecting the final model, with all QTL and epistasis, was the Akaike information criteria (AIC) (Akaike 1974):

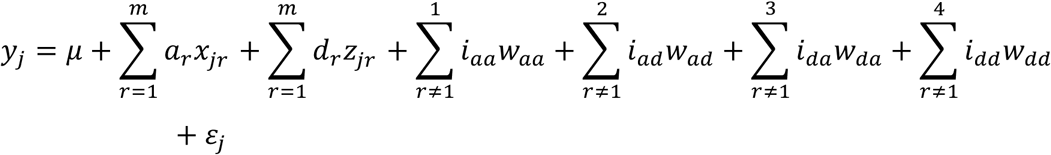

Where: *y*_j_ is the adjusted phenotypic mean for the 𝑗 individual (j = 1, 2,…, 200); 𝜇 is the intercept; 𝑎_𝑟_ the additive effect of QTL 𝑟 ; 𝑑_𝑟_ the dominance effect of QTL 𝑟, and *i*_𝑎𝑎_, *i*_𝑎𝑑_, *i*_𝑑𝑎_, and *i*_𝑑𝑑_ the epistatic effects additive x additive, additive x dominance, dominance x additive and dominance x dominance, respectively. 𝑥_j𝑟_ is an indicator variable according to Cockerham’s genetic model (Cockerham 1954). For each progeny, the explanatory variable 𝑥_j𝑟_ may assume values equal to 1, 0 or -1, if the QTL is QrQr, Qrqr, or qrqr, respectively; variable 𝑧_j𝑟_ assumes values of ½ if Qr is QrQr or -1/2, in other cases. The model accounts for epistasis, where the regression coefficients for epistatic effects represents 𝑤_𝑎𝑎_ = 𝑥_j𝑟_ × 𝑥_j𝑠_; 𝑤_𝑎𝑑_ = 𝑥_j𝑟_ × 𝑧_j𝑠_; 𝑤_𝑑𝑎_ = 𝑧_j𝑟_ × 𝑥_j𝑠_; and 𝑤_𝑑𝑑_ = 𝑧_j𝑟_ × 𝑧_j𝑠_ between QTL *r* and *s*, and 𝜀_j_ represents the residues assumed to be *N*(0, *σ*^2^).

For marker-assisted selection (MAS), we computed the probabilities of individuals harboring favorable QTL genotypes, based on the MIM model and on the genetic map. Then, for the final QTL model, we estimated the fitted *y*^ values and the residuals, from the difference of observed (*y*) and fitted values (*y*^_𝜄_ = *y*_*i*_ − residual_*i*_*y*_*i*_residual_*i*_). The fitted *y*^ values based on the model were used to rank the individuals, and better genotypes were found among the most valuable ones. The correlation between *y*^ and *y* (observed values) was used to estimate the prediction accuracy. This gave us insights into the model’s predictive ability and reliability in guiding the selection of individuals with enhanced resistance to RKN. QTL analysis was carried out with the R/qtl package (Broman et al. 2003).

### Characterization of genomic region and discovery of candidate genes

The genomic context of the QTL was investigated to identify candidate resistance genes. SNP flanking markers were defined based on the highest LOD score values derived from scanone analysis using the MIM model. The reference genome of Phaseolus vulgaris v2.1 (Schmutz et al. 2014), available on the Phytozome database (https://phytozome-next.jgi.doe.gov/), provided the protein sequence annotations for the coding genes within the genomic interval. The functional domains and predicted protein functions were analyzed using the online databases InterPro (https://www.ebi.ac.uk/interpro/) and Uniprot (https://www.uniprot.org/) (accessed on 2 December 2024), respectively. Additionally, candidate genes were identified among those reported by Orsi et al. (2024) as differentially expressed during the *P. vulgaris*–*M. incognita* interaction by RNA-sequencing analysis.

## Results

### Phenotypical performance of the TB population

In this study, 200 F_2:3_ families, along with the parents and two checks (resistant and susceptible) were evaluated. The population exhibited high phenotypic variation for the three measured traits: egg mass (EM), root-galling index (GI) and root dry mass (RM). The adjusted means of the traits followed continuous distributions and indicated the absence of extreme outliers. The adjusted means for EM were 92.9 for IAC-Tybatã and 221.08 for Branquinho, while for IG, the adjusted means were 1.52 and 2.208, respectively. The number of EM ranged from 52 to 308 across the population, with an average of 163.5, GI had an average of 2.11, ranging from 1.3 to 2.9, and RM varied from 44 mg to 100 mg, with an average of 75 mg (Supplementary Figure 1). The heritability was 78.1% for EM, 82.4% for GI, and 67.8% for RM, suggesting high odds of finding QTL. The correlation coefficients (Supplementary Figure 2) showed a high correlation between EM and GI (0.81). Additionally, RM correlated with EM (0.75) and GI (0.68) (Supplemental Figure 2).

### 11 linkage groups for QTL detection

A linkage map was constructed for the *F*_2_ T x B population with 954 SNPs allocated in 11 linkage groups, totaling 1,687 cM (Figure 1). Markers were aligned with the published reference genome, allowing assignment of the linkage groups to chromosomes (Supplementary figure 4). Subsequently, examination of the heatmaps (Supplementary figure 3) facilitated removal of unlinked markers from the map. Importantly, all eleven linkage groups exhibited a consistent alignment with the reference genome. This alignment was validated against a global error rate probability of 5% (global error 0.05), as outlined by Taniguti et al. (2022). Each linkage group had around 90-100 markers, with inter-marker distances ranging from 5 cM to 25 cM. Gaps were observed around the centromere regions, probably due to hypermethylated areas, as this is a feature of GBS markers (Supplementary figure 4).

**Figure 1:**
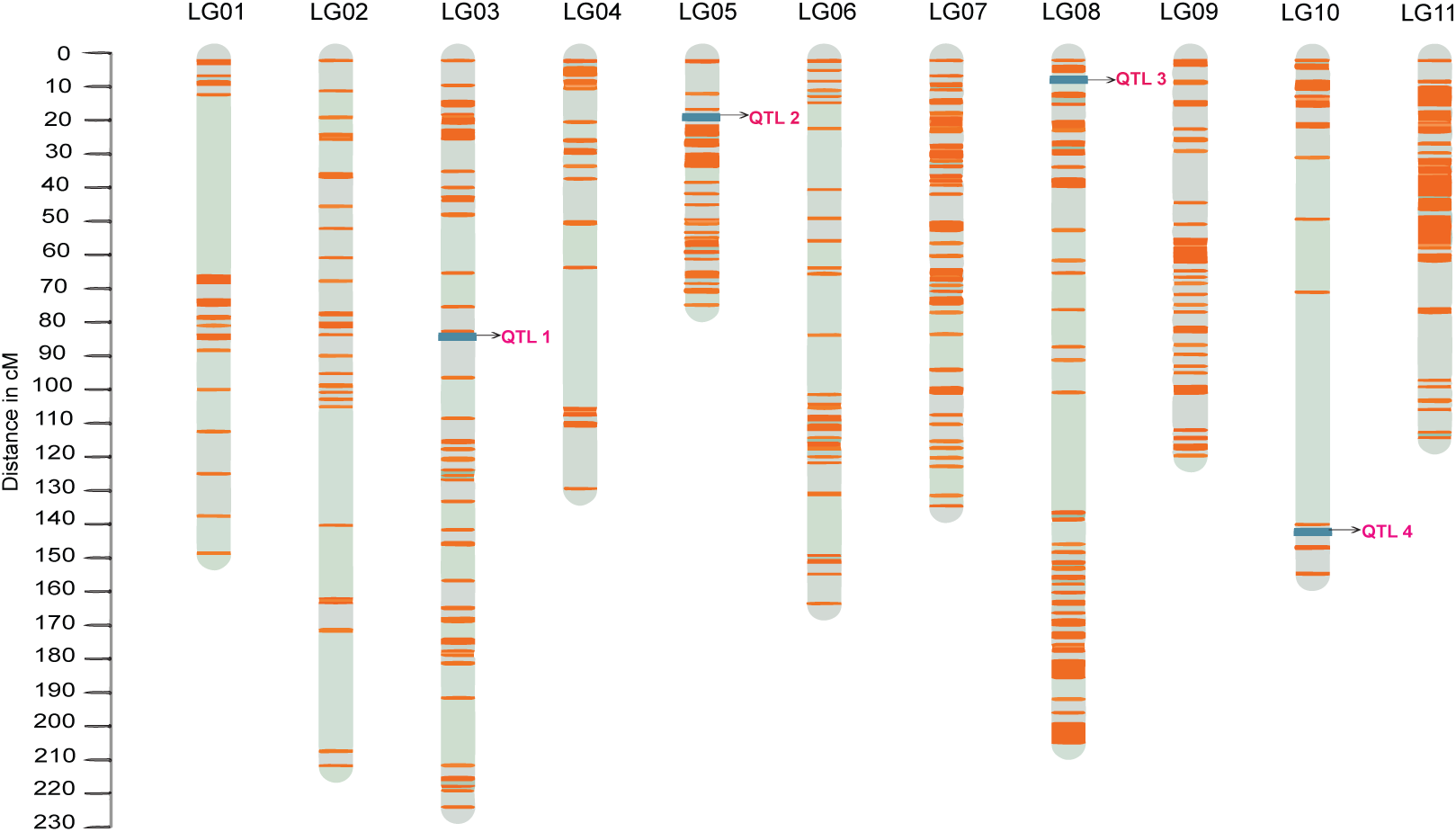
Linkage Map with Identified QTLs. The orange lines represent genetic markers, and the pink lines indicate the position of significant QTLs. The map covers LG01 to LG11, with QTL 1 mapped on LG03, QTL 2 on LG05, QTL 3 on LG08, and QTL 4 on LG10. The y-axis indicates the genetic distance in centiMorgans (cM).

### QTL identification for RKN disease

Subsequent investigation focused on identifying significant QTL associated with the host response to *M. incognita* inoculation, specifically for the RKN resistance traits egg mass (EM), root-galling index (GI), and root mass (RM). During initial analysis using interval mapping (IM) (Figure 2a), significant results were observed only for GI, with a LOD peak of 3.7 on chromosome 10. For composite interval mapping (CIM), the LOD score for this QTL increased to 6.30, as expected, as this model has more statistical power, due to the incorporation of markers as covariates (Figure 2b). No significant QTLs were identified for EM and RM, so no further analysis was carried out for these traits.

**Figure 2:**
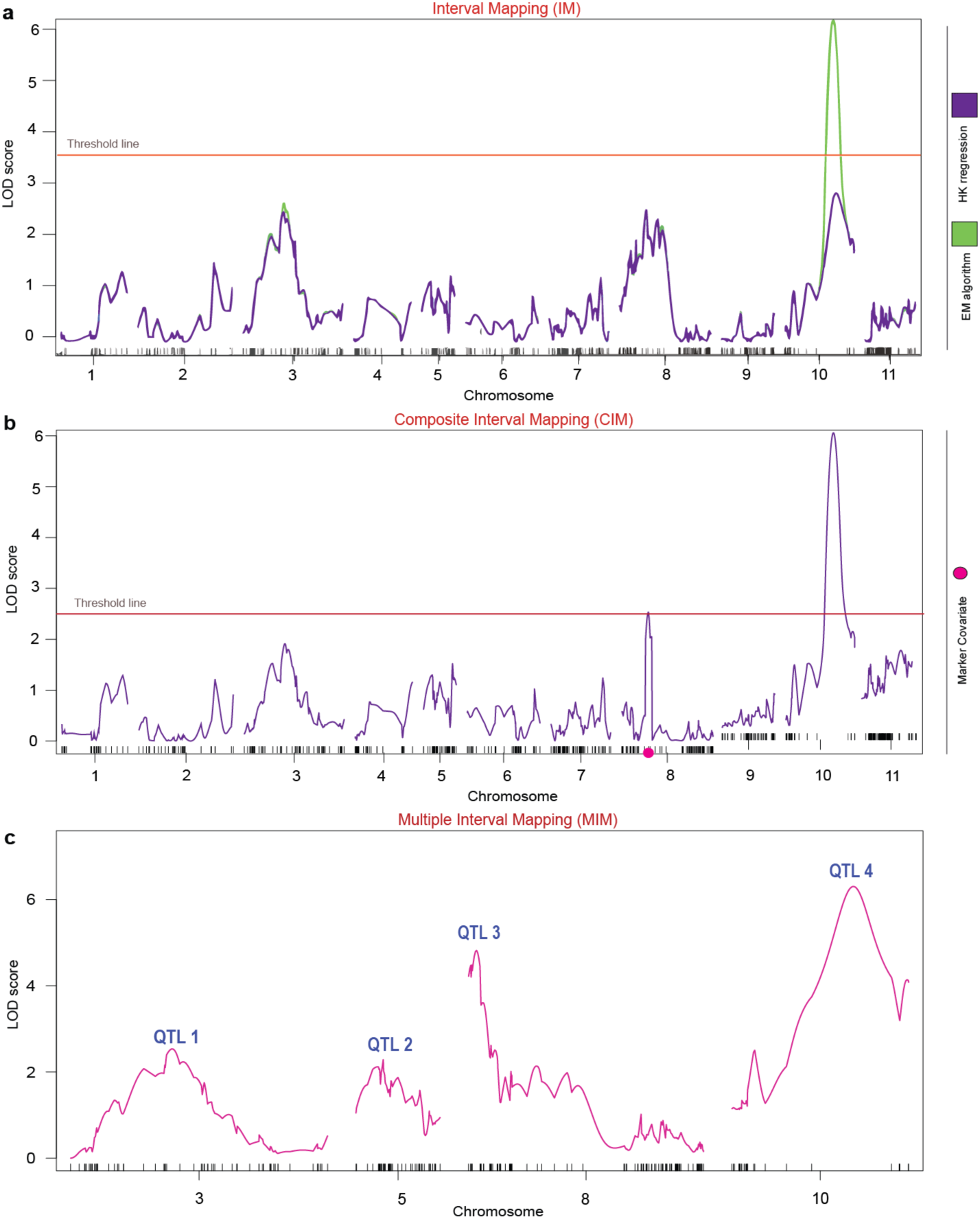
Identification of Quantitative Trait Loci (QTL) for Root-galling index (GI) using different statistical models. (a) **Interval Mapping (IM)**: The LOD scores for the gall index (GI) are plotted across the genome. A significant QTL peak is identified on Pv10, exceeding the threshold line set at LOD = 3.7, providing statistical evidence of a QTL associated with the trait. (b) **Composite Interval Mapping (CIM)**: This method refines the QTL analysis by including co-factors to control for genetic background noise. The presence of the previous QTL was confirmed with a higher LOD Score (6.30). (c) **Multiple Interval Mapping (MIM)**: The final QTL model identifies four significant QTLs (numbered 1 to 4) for GI, located on chromosomes Pv03, Pv05, Pv08, and Pv10. The MIM model also allowed testing for epistatic interactions, enhancing the precision of QTL detection and providing comprehensive estimation of the genetic architecture of the trait.

**Figure 3:**
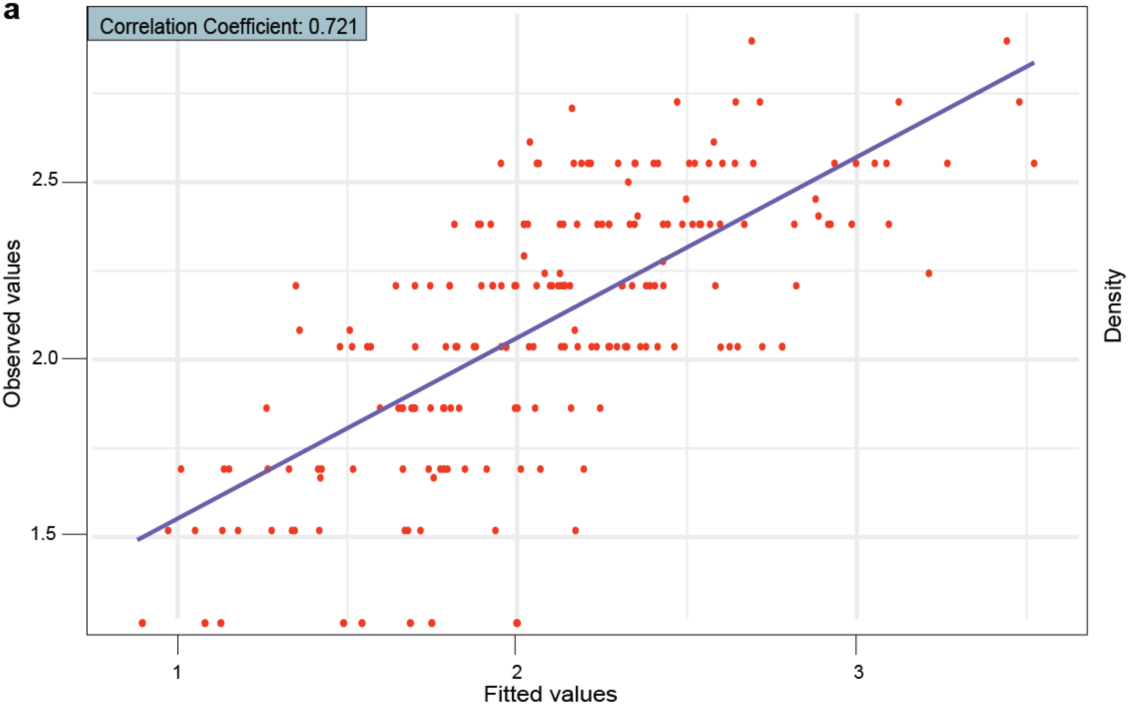
Scatter plot comparing genotypic values predicted from the selected QTL model and actual values for root-galling index (GI). A correlation coefficient of 0.72 was observed.

### Multiple interval mapping shows QTL on four chromosomes related to GI

Based on the results of the CIM analysis, we proceeded with multiple interval mapping (MIM), starting with the QTL identified on chromosome Pv10. Evidence of QTLs was found on chromosomes Pv03, Pv05, and Pv08 (Figures 1 and 2). A test for two-way epistatic interactions was then carried out and significant interaction between QTLs identified on chromosomes Pv08 and Pv10 (Table 2). The final model explained 22.44% of the phenotypic variance (%PVE), with a total LOD score of 11.04, and the model was statistically significant at p-value = 0.05 (Table 1). In an F_2_ population, R/qtl assigns 1 degree of freedom (*df*) for the additive effect of each QTL and 1 df for the dominance effect, resulting in 2 *df* per QTL. Since our model contains four QTLs, this accounts for 8 *df*. Additionally, the interaction between QTLs yields 4 *df*, as both QTLs have additive and dominance effects interacting. This resulted in a total of 12 *df* for the ANOVA table (Table 1). The results were extracted with the function summary from *fitqtl*.

**Table 1:** Analysis of variance (ANOVA) from the MIM (Multiple Interval Mapping) model. The model has 12 degrees of freedom, corresponding to all major effects (additive and dominance) of each QTL plus 4 epistatic interactions between QTL 3 and 4.

| Term | df | SS | MS | LOD | % PVE | P-value |
| --- | --- | --- | --- | --- | --- | --- |
| **Model** | 12 | 5.950 | 0.496 | 11.04 | 22.44 | 2.30e-06 |
| **Error** | 187 | 20.565 | 0.110 | - | - | - |
| **Total** | 199 | 26.516 | - | - | - | - |

**Table 2:** “Drop one QTL at time” by MIM for the Root-galling index (GI)

| QTL | df | LOD | % PVE | P-value |
| --- | --- | --- | --- | --- |
| QTL1: 89.0 | 2 | 2.54 | 4.66 | 0.0043 |
| QTL2: 24.0 | 2 | 2.29 | 4.19 | 0.0073 |
| QTL3: 7.0 | 6 | 4.82 | 9.10 | 0.0019 |
| QTL4: 107.0 | 6 | 6.30 | 12.12 | 0.0001 |
| QTL3: 7.0 × QTL4: 107.0 | 4 | 4.16 | 7.79 | 0.0012 |

*QTL* indicates the chromosome and position where the *QTL* is located. *Df* stands for degrees of freedom. *LOD*: Logarithm of the odds score indicating the strength of the association. *%PVE*: Percentage of phenotypic variance explained by the *QTL*. *P-value* Statistical significance of the *QTL*. The degrees of freedom for *QTL* 3 and 4 include the epistatic interactions.

The function *fitqtl* also includes the “drop one QTL at a time” table, where each QTL is evaluated separately by the algorithm. Each locus contributes 2 *df*, except for QTL on chromosome 8 and chromosome 10, which contribute 6 *df,* as they have significant epistatic interactions. The QTL accounted for 4%, 5%, 9%, and 12% of the phenotypic variation on chromosomes Pv03, Pv05, Pv08, and Pv10, respectively. Additionally, the interaction between the QTL on chromosomes Pv08 and Pv10 yielded 8% of the phenotypic variation (Table 2). The interactions between the additive and dominance effects of these QTL are marginally significant. These findings are summarized in Table 3, where we detail each effect, while considering the effects of *a; d;* 𝑎 × 𝑎,𝑑 × 𝑎, 𝑎 × 𝑑 *and* 𝑑 × 𝑑.

**Table 3:** Estimated effects of QTLs and their interactions.

| Parameter | Estimate<br>(est) | Standard Error<br>(SE) | t Value |
| --- | --- | --- | --- |
| *Intercept* | 2.090 | 0.026 | 80.10 |
| QTL 1: 89.0 (*a*) | -0.089 | 0.033 | -2.67 |
| QTL 1: 89.0 (*d*) | -0.115 | 0.056 | -2.05 |
| QTL 2: 24.0 (*a*) | -0.109 | 0.035 | -3.08 |
| QTL 2: 24.0 (*d*) | 0.031 | 0.049 | 0.63 |
| QTL 3: 7.0 (*a*) | -0.066 | 0.037 | -1.75 |
| QTL 3: 7.0 (*d*) | -0.049 | 0.054 | -0.90 |
| QTL 4: 107.0 (*a*) | 0.183 | 0.053 | 3.42 |
| QTL 4: 107.0 (*d*) | 0.223 | 0.155 | 1.44 |
| QTL 3: 7.0 × QTL 4: 107.0<br>(*a* × *a*) | 0.195 | 0.085 | 2.30 |
| QTL 3: 7.0 × QTL 4: 107.0<br>(*d* × *a*) | -0.411 | 0.115 | -3.56 |
| QTL 3: 7.0 × QTL 4: 107.0<br>(*a* × *d*) | -0.021 | 0.238 | -0.09 |
| QTL 3: 7.0 × QTL 4: 107.0<br>(*d* × *d*) | -0.013 | 0.331 | -0.04 |

The notations (*a*) and (*d*) represent additive and dominance effects, respectively. The epistatic interactions between QTLs are of type *a*×*a*, *a* × *d*, *d* × *a* and *d* × *d*. The number following the QTL, represents the map position of the QTL on the chromosome.

Using the same QTLs identified based on MIM, genotypic values were predicted for 200 individuals from the F_2:3_ segregating population related to the target trait, GI. The fitted values (*y*^) for each F_2:3_ individual was calculated by subtracting the residuals from the true phenotypes. These fitted values represent the portion of the trait explained by the QTL. High resistance to gall formation was identified in several individuals, as they had genotypes that combined favorable alleles at the significant QTL. These geno-typic values were used to select individuals for future breeding programs, aimed at enhancing resistance traits by leveraging the genetic information provided by the MAS approach. The correlation between the observed and fitted values was high (0.72), indicating promising model predictivity for marker assisted selection, suggesting that the model is performing fairly well (Figure 4).

### Candidate gene search

The search for candidate genes focused on the quantitative locus located on chromosome Pv10, which exhibited the highest LOD score across the three analyses (IM, CIM, and MIM), and demonstrated the greatest consistency. The candidate gene interval, defined by the SNP markers S10_42850891 and S10_43584857, spans 733,966 bp and 102 genes (excluding isoforms), from *Phvul.010G147100* to *Phvul.010G157100*. Of these, 23 genes correspond to findings reported by Orsi et al. (2024) (Supplementary Table S2), highlighting their relevance as genes for further investigation. Additionally, the genes *Phvul.010G152200*, *Phvul.010G154000*, and *Phvul.010G155800* encode proteins with domains previously associated with nematode response (Table 4). This targeted selection represents a strategic approach, to further explore and validate the possible functional significance of these specific genes’ mechanisms of resistance and susceptibility to the studied trait, including gene editing.

**Table 4:**
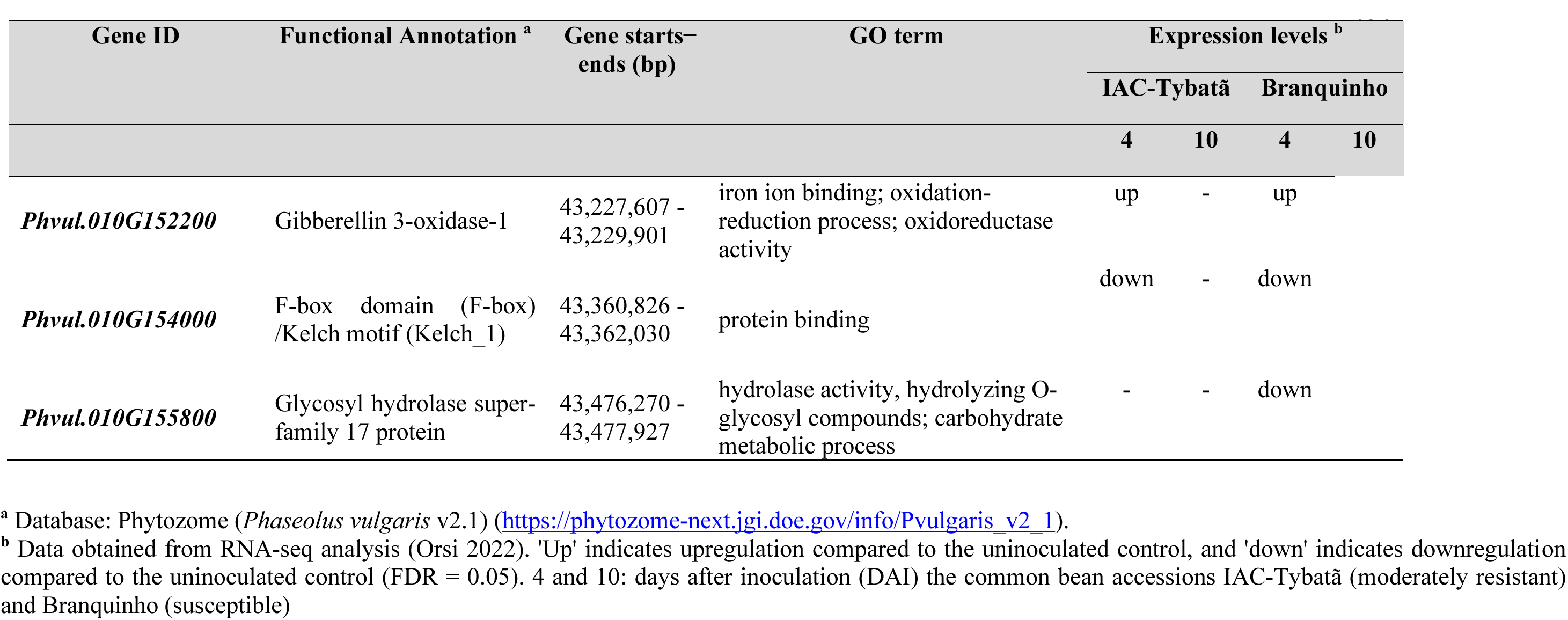
Functional annotation of genes associated with plant-nematode interactions on chromosome 10 of *Phaseolus vulgaris*, within the flanking marker interval of QTL 4 on Pv10.

## Discussion

The parental line responses to RKN were consistent with that reported in a diversity panel evaluation of the IAC germplasm (Giordani et al. 2022), demonstrating the viability of the reported phenotyping protocol and the reliability of the experiments. The high heritability estimates for GI (82.4%), EM (78.1%), and RM (67.8%) suggest these traits are predominantly influenced by genetic factors. However, a significant QTL was identified only for GI. Although the phenotypic analysis showed a strong correlation between EM and GI (0.81), the twofold phenotypic contrast between the parents in the number of egg masses was insufficient to identify a QTL associated with the EM trait. Increasing the number of individuals evaluated in the population could improve the chances of identifying a QTL linked to this trait.

While some studies have explored root-knot nematode resistance in common beans, identifying significant QTL and candidate genes associated with resistance, further investigation into its genetic architecture through QTL mapping is essential. A key step in this process is to construct a high-quality linkage map, critical for reliable QTL identification. Several factors may contribute to the presence of unlinked markers or gaps in linkage maps (Supplementary Figures 3 and 4). These challenges can arise at various stages of the process, including GBS library construction, the influence of hypermethylated regions near centromeres, DNA sequencing, and bioinformatics analyses applied to the dataset. To mitigate these issues, specialized tools like Onemap software (Margarido et al. 2007) offer refined methods of map construction, significantly improving the accuracy and completeness of linkage maps. Plots depicting correlation between the reference genome and marker distances within linkage groups (Supplementary Figure 2) provide a visual assessment of the consistency and alignment of the constructed map with the genome. Additionally, efforts were made to ensure that the final map size aligns with established findings, as reported in studies by Giordani et al. (2022) and Bassett (1988).

Reliable statistical analyses were performed based on phenotypic estimates. The CIM method provided a significant advantage, using covariate markers as boundaries for flanking markers, which enabled precise QTL localization (Liu 1998). This provided a robust foundation for fitting the MIM model in the next step. The final model, refined through the MIM method, identified significant QTL associated with GI and revealed epistatic interactions. This approach enhances our understanding of the genetic architecture basis of resistance. Resistance to RKN, measured by GI, was found to be a quantitative trait controlled by a few QTL with moderate to large effects, some of which exhibited significant epistatic interactions.

The fitted model provided valuable insights into the genetic basis of nematode resistance, specifically regarding the parental contributions of IAC-Tybatã and Branquinho, referred to here as moderately resistant (MR) and susceptible (S) lines, respectively. The identified QTL accounted for a considerable proportion of the additive and dominance variance associated with resistance traits. The small percentage of phenotypic variance explained (PVE) in response to RKN aligns with expectations for complex traits, where PVE reflects the heritability of each identified QTL. This limited PVE is commonly observed in nematode disease-related traits in plants, highlighting the intricate genetic architecture underlying them. Similar patterns of small PVE have been reported in various crops for nematode resistance. Giordani et al. (2022) observed a comparable RKN resistance trend in the common bean. In studies of other crops, such as rice (*Oryza sativa*), in response to the root-knot nematode *Meloidogyne graminicola* (Galeng-Lawilao et al. 2020), soybean (*Glycine max*), in response to the Soybean Cyst Nematode (SCN, *Heterodera glycines*) (Huang et al. 2021), and peanut (*Arachis hypogaea* L.), in relation to the root-knot nematode *M. arenaria* (Burow et al. 2014), similar challenges have been encountered in explaining a significant proportion of phenotypic variance.

The MIM model identified four QTLs associated with GI: on chromosomes Pv03 (QTL1), Pv05 (QTL2), Pv08 (QTL3), and Pv10 (QTL4). For QTLs 1 and 2, the negative additive effect indicates that alleles from the moderately resistant line contribute to increased resistance (Table 3). This finding suggests that selecting for these alleles may enhance resistance in future breeding programs. Targeted selection for favorable alleles at QTL 2 and QTL 4 could accelerate breeding efforts to improve nematode resistance in subsequent populations. Interestingly, QTL 4 exhibited the opposite effect, as alleles from the susceptible parent increased the trait value, contributing to greater susceptibility. This underscores the critical role of this locus in terms of susceptibility. Selection against the allele from the S line may reduce the susceptibility through marker assisted selection.

For QTL 2 at 24 cM (Figure 1), our findings are consistent with those of Giordani et al. (2022). In their study, using a BC_2_F_4_ population inoculated with *M. incognita* race 3, a significant QTL associated with GI was identified at the extremity of Chr05, at 18.552 cM. While the physical locations do not overlap, the relatively small distance in cM suggests a potential connection. Additionally, GWAS analysis of the IAC germplasm panel identified a significant SNP located at the end of Pv05 (35,206,731 bp), which closely corresponds to the Chr05 endpoint (32,068,861 bp) in the present study. Notably, the QTL 3 is located near a marker at 8,890,077 bp, which aligns with the *Phvul.008G089600* locus (8,992,049 to 8,996,924 bp) associated with SCN resistance in the common bean, as reported by Shi et al. (2021). Moreover, Jain et al. (2019) identified homologs of the soybean *Rhg1* locus on Pv01 and Pv08. These were found in both the Middle American and Andean gene pools of the common bean. The close genomic proximity and supporting evidence from previous studies highlight the importance of this region on Pv08 in conferring resistance to nematodes.

To further elucidate the genetic basis of QTL 4 on Pv10, the interval with the highest LOD scores, defined by the MIM analysis, was used to identify candidate genes associated with the GI trait. Building on these findings, Giordani et al. (2022) identified 14 resistance gene analogs (RGA) associated with GI and EM traits by evaluating a panel which included IAC-Tybatã and Branquinho genotypes, both inoculated with *M. incognita*. Additionally, Orsi et al. (2024) characterized genes involved in the common bean response to *M. incognita*, focusing on those differentially expressed between the same MR and S genotypes. Remarkably, RNA-seq analysis identified 23 out of 102 genes located within the QTL 4 interval.

The genes identified in our study encompass a diverse range of significant functions, including three key genes within the QTL 4 interval previously linked to plant-nematode interactions. Among these, the *Phvul.010G152200* gene is involved in hormonal pathways, particularly in the biosynthesis and signaling of gibberellins. These hormones, along with cytokinins, are known to facilitate the formation of nematode feeding sites, playing a crucial role in establishing parasitic relationships (Siddique and Grundler, 2018). Supporting this, a study demonstrated that *M. graminicola* infection leads to gibberellin accumulation at the infection site in rice (Yimer et al., 2018). Their findings revealed that gibberellin influences RKN parasitism in rice by antagonizing jasmonate-induced defenses. This is consistent with the observed upregulation of the gibberellin-related gene *Phvul.010G152200* in both MR and S genotypes at 4 days after inoculation (DAI), during the early stages of *M. incognita* infection establishment (Table 4).

The other gene model, *Phvul.010G154000*, encodes an F-box/Kelch-Repeat protein previously associated with nematode susceptibility. It has been shown that the Fbox protein (At2g44130) from *Arabidopsis thaliana* is induced by *M. incognita* during the early stages of feeding site formation (Curtis et al., 2013). Additionally, overexpression of this protein resulted in a 67% increase in nematode infection, potentially due to enhanced attraction of *M. incognita* juveniles (J2) to root exudates. The authors hypothesized that At2g44130 functions as a susceptibility gene for nematode infection. The downregulation of *Phvul.010G154000* observed in both MR and S genotypes at 4 DAI (Orsi et al. 2024) may represent an early attempt to block nematode infection. These genotypes primarily differ in their transcriptomic profiles during gall formation at later stages, notably at 10 DAI.

Lastly, the gene *Phvul.010G155800* belongs to the glycosyl hydrolase superfamily 17, a group which encodes proteins previously associated with plant defense mechanisms. A similar protein in *A. thaliana* (*At4g16260*) seemingly plays a role in defense against the beet cyst nematode *Heterodera schachtii* (Hamamouch et al., 2012). At4g16260 encodes a putative beta-1,3-endoglucanase from the glycosyl superfamily, considered a pathogenesis related protein that interacts with the 30C02 cyst nematode effector. Plants overexpressing *At4g16260* exhibited reduced nematode infection, suggesting *H. schachtii* manipulates the plant by reducing the expression of this gene in feeding sites to promote successful parasitism (Hamamouch et al., 2012). Similarly, the downregulation of *Phvul.010G155800* observed in Branquinho genotype at 4 DAI (Orsi et al. 2024) indicates a similar manipulation by the nematode to suppress the plant’s defense. However, the complex relationships among these candidate genes and their specific roles in conferring resistance require further investigation. Ongoing research into the interplay of these genes will provide deeper insights into the molecular mechanisms underlying RKN resistance in common beans.

The additive × additive epistatic interaction between QTL 3 and QTL 4 suggests that combining favorable alleles from both loci confers greater resistance than either allele alone. However, the dominance × additive interaction reveals a more complex relationship, where heterozygous alleles at QTL 4, when combined with homozygous alleles at QTL 3, reduce the trait value. This interaction reflects the challenges of working with populations that include heterozygotes, as such interactions can be difficult to manage in a breeding context. Moreover, this complexity is particularly relevant in self-pollinating species, where epistatic interactions between QTLs, especially additive × additive, can significantly influence the heterosis (Garcia et al. 2008). Supporting this, research in other plants, such as the study by Dodia et al. (2019) on stem rot resistance in cultivated peanut and Den Boer et al. (2014) on downy mildew resistance in lettuce, demonstrate the essential role of epistatic interactions in controlling disease resistance.

In conclusion, the TB population, reflecting the contrast between the parents, is a valuable resource for investigating the genetic architecture of resistance to RKN in the F_2_ generation. The linkage map successfully integrated 954 SNPs into 11 linkage groups spanning 1,687 cM and provided a solid foundation for the IM and CIM approaches, which were further refined through MIM analysis. This model identified four significant QTLs on chromosomes Pv03, Pv05, Pv08, and Pv10, along with epistatic interactions. The MIM model also allowed calculation of genotypic values, enabling marker-assisted selection (MAS). The high correlation between predicted and observed values (0.72) attests to the model’s accuracy and relevance. This study demonstrates the importance of combining genomic, bioinformatic, and statistical genetics approaches to enhance resistance to RKN in common bean cultivars. These multidisciplinary strategies are crucial for breeders, providing effective tools for utilizing QTL data to improve the efficiency of breeding programs and develop pathogen-resistant varieties.

## Data availability

All experimental data are included in the manuscript or are available in supplementary materials.

## Author contributions

T. O. Floriani: Formal analysis, methodology, investigation, writing: original draft, review and editing. H. C. Gama: Investigation, formal analysis, methodology. B. M. Moreno: Methodology. G. A. L. Costa: Formal analysis. W. Giordani: Methodology. A. F. Chiorato: Resources. L. B. J. Bibiano: Writing: review and editing. T. A. Parker: Methodology, writing: review. L. E. A. Camargo: Investigation, writing: review and editing, M. L. Carneiro Vieira: Conceptualization, funding acquisition, resources, writing: review and editing, A. A. F. Garcia: Supervision, investigation, funding acquisition, resources, writing: review and editing.

## Acknowledgments

We would like to thank Carlos A. de Oliveira for excellent technical assistance, and Professor Mario Massayuki Inomoto (University of São Paulo) for providing the nematode inoculum.

## Funding

This research was supported by the following Brazilian Institutions: Fundação de Amparo à Pesquisa do Estado de São Paulo (FAPESP, Grant no. 2020/02755-2 and 2022/04061-3), Conselho Nacional de Desenvolvimento Científico e Tecnológico (CNPq, Grants no. 131678/2021-3 and 313269/2021-1), Coordenação de Aperfeiçoamento de Pessoal de Ensino Superior (CAPES, Finance Code 001).

## Conflicts of interest

The authors declare no conflict of interest.

---

**Supplementary Figure 1:**
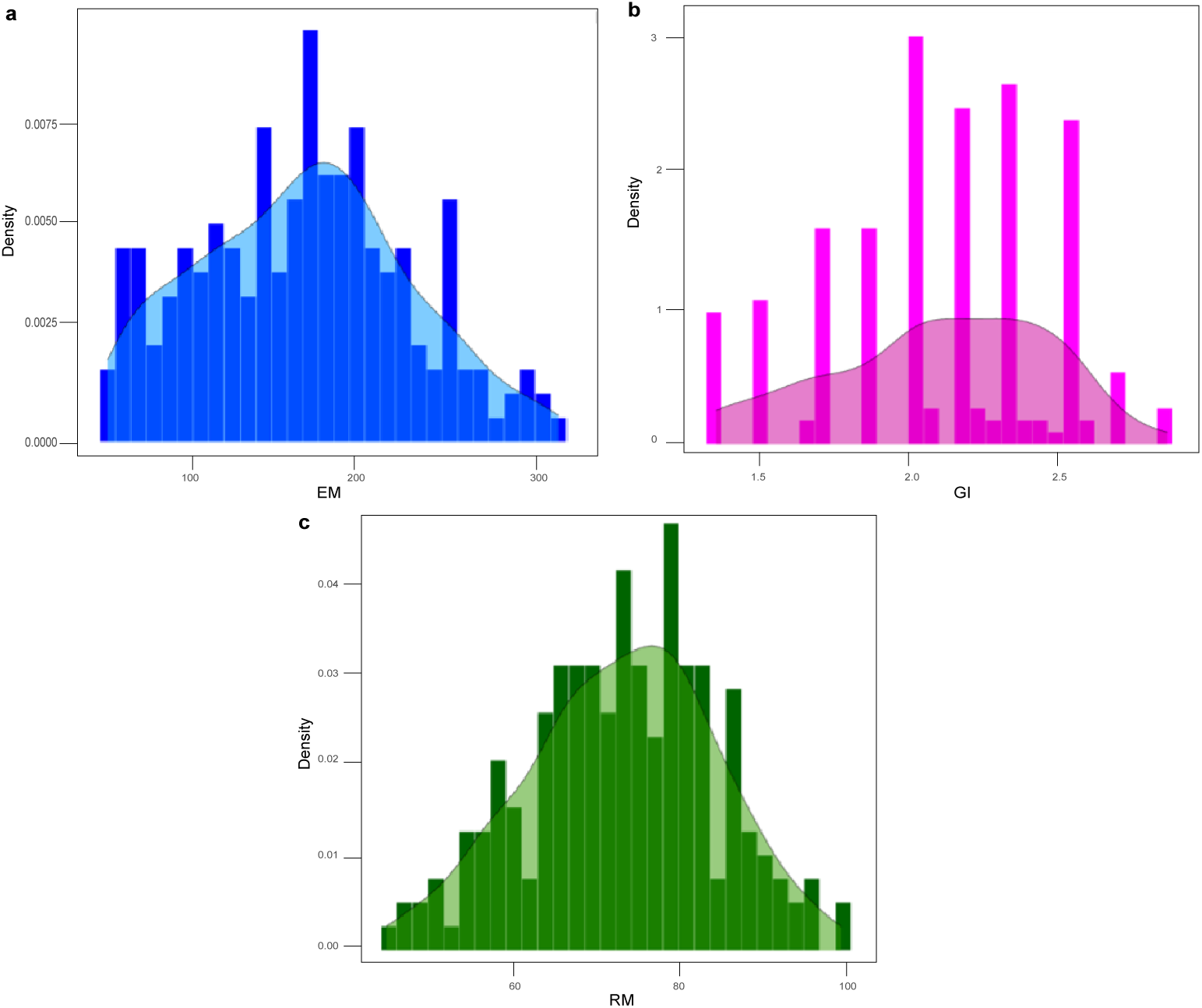
Frequency distribution of phenotypic traits in the F_2:3_ families. **(a)** Egg Mass (EM) **(b)** Root-Galling Index (GI)**(c)** Root Dry Mass (RM)

**Supplementary Figure 2:**
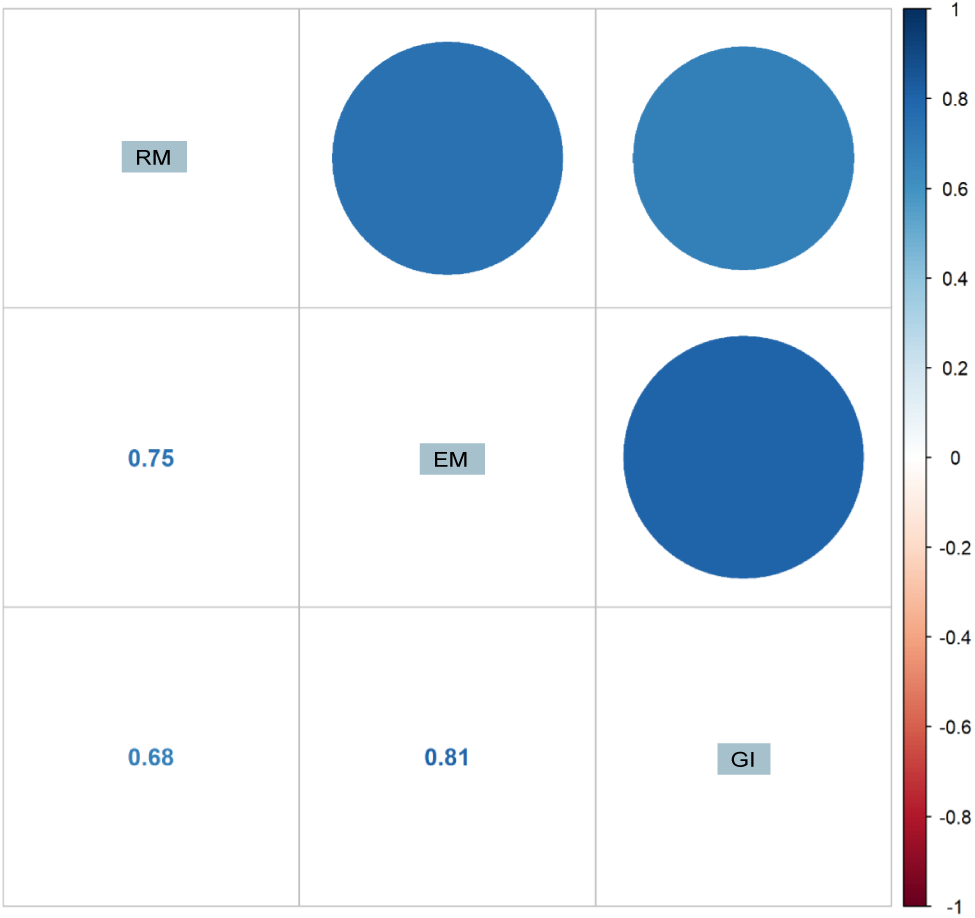
Correlation matrix of phenotypic traits showing the relationships between root mass (RM), egg mass (EM), and gall index (GI). The circle size represents the strength of the correlation, with numerical values indicating correlation coefficients.

**Supplementary Figure 3:**
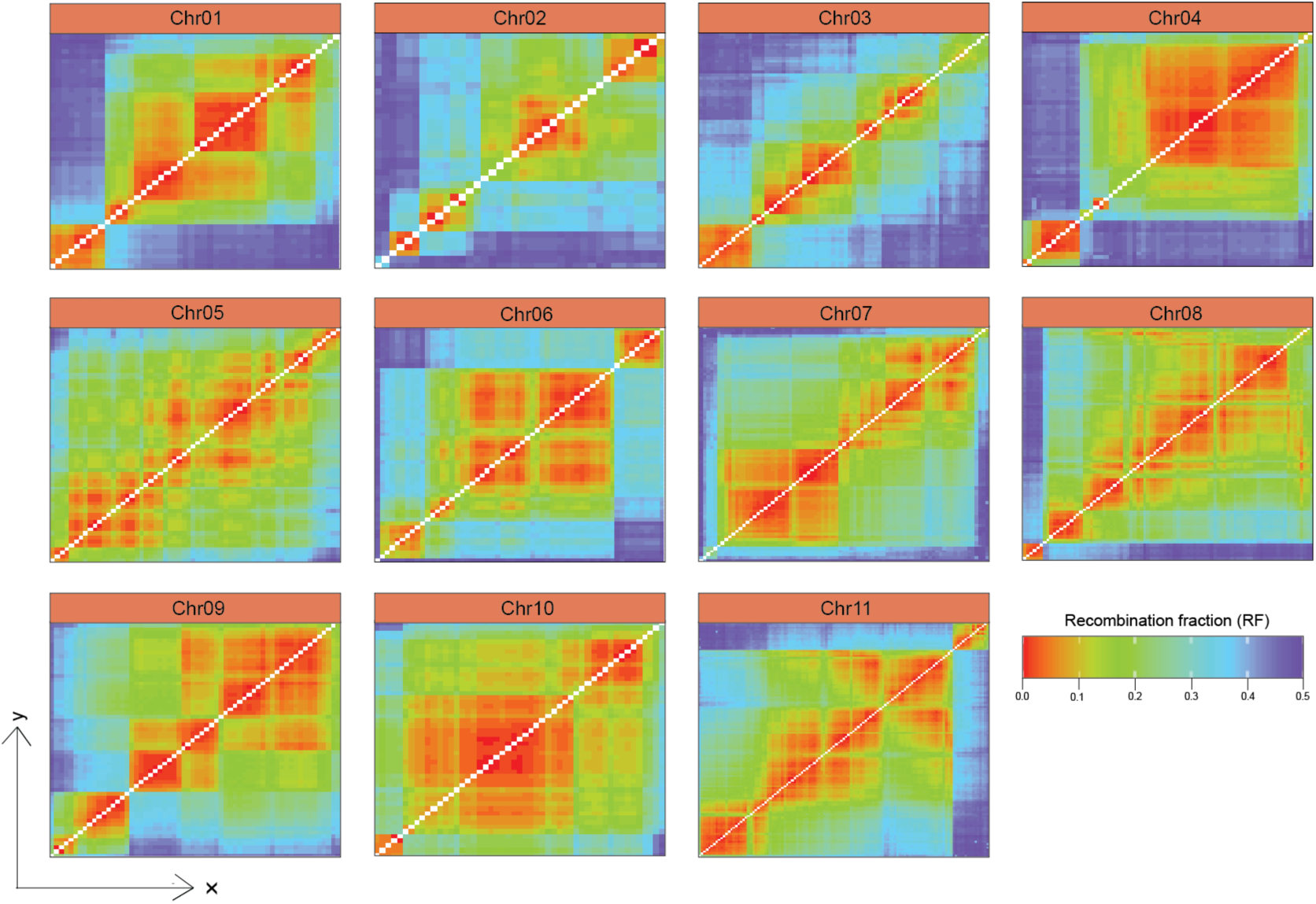
Heatmaps displaying the recombination fraction between markers in 11 chromosomes.

**Supplementary Figure 4:**
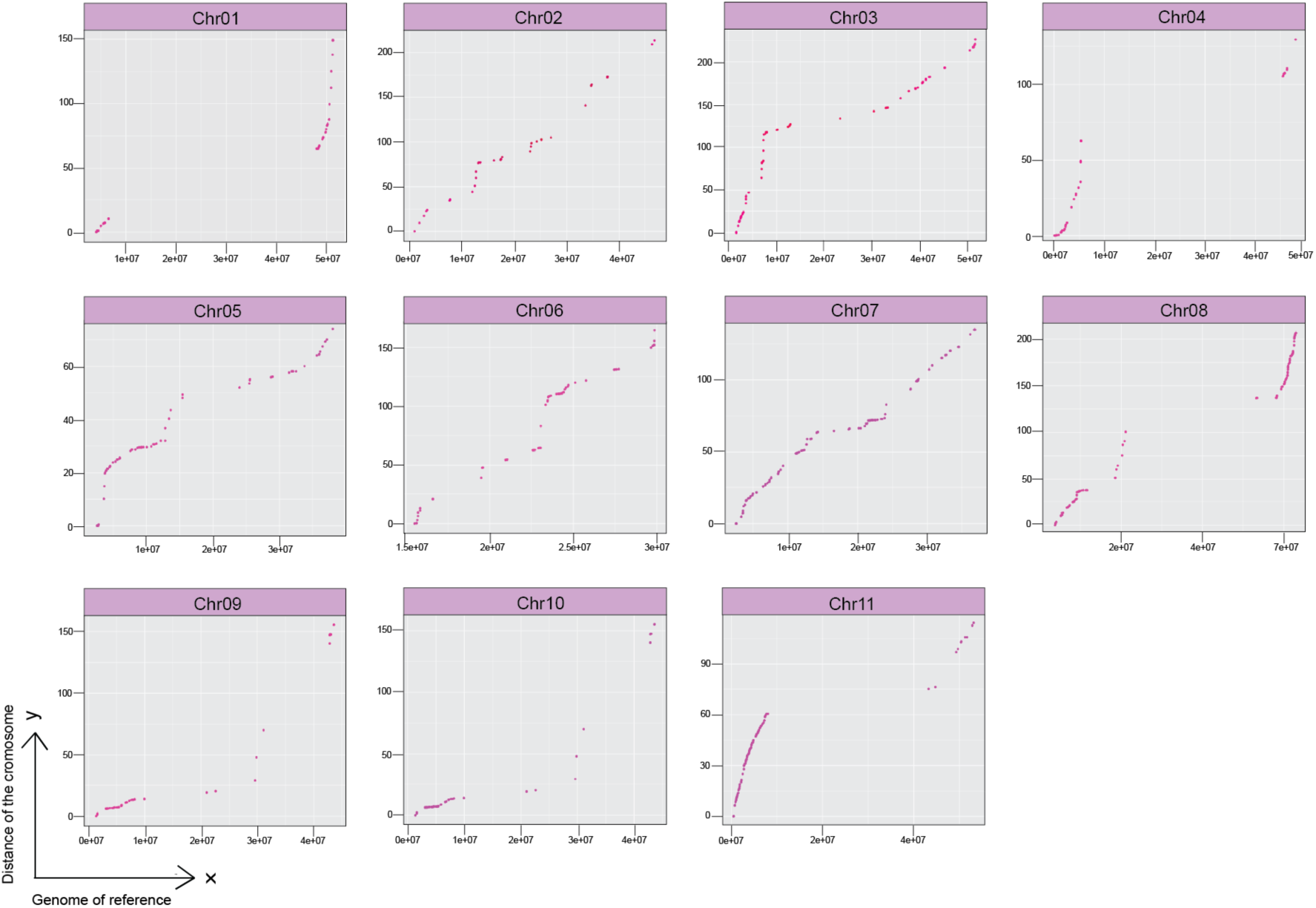
Relationship between the genetic map and the common bean reference genome. The scatter plots depict the relationship between the physical positions of markers on the reference genome (x-axis) and their corresponding genetic map positions in centiMorgans (y-axis) for each chromosome (Pv01 to Pv11). Each point represents a marker, and the alignment along the diagonal indicates the consistency of the genetic map with the reference genome. Gaps show a possible centromere region.

**Supplementary Figure 5:**
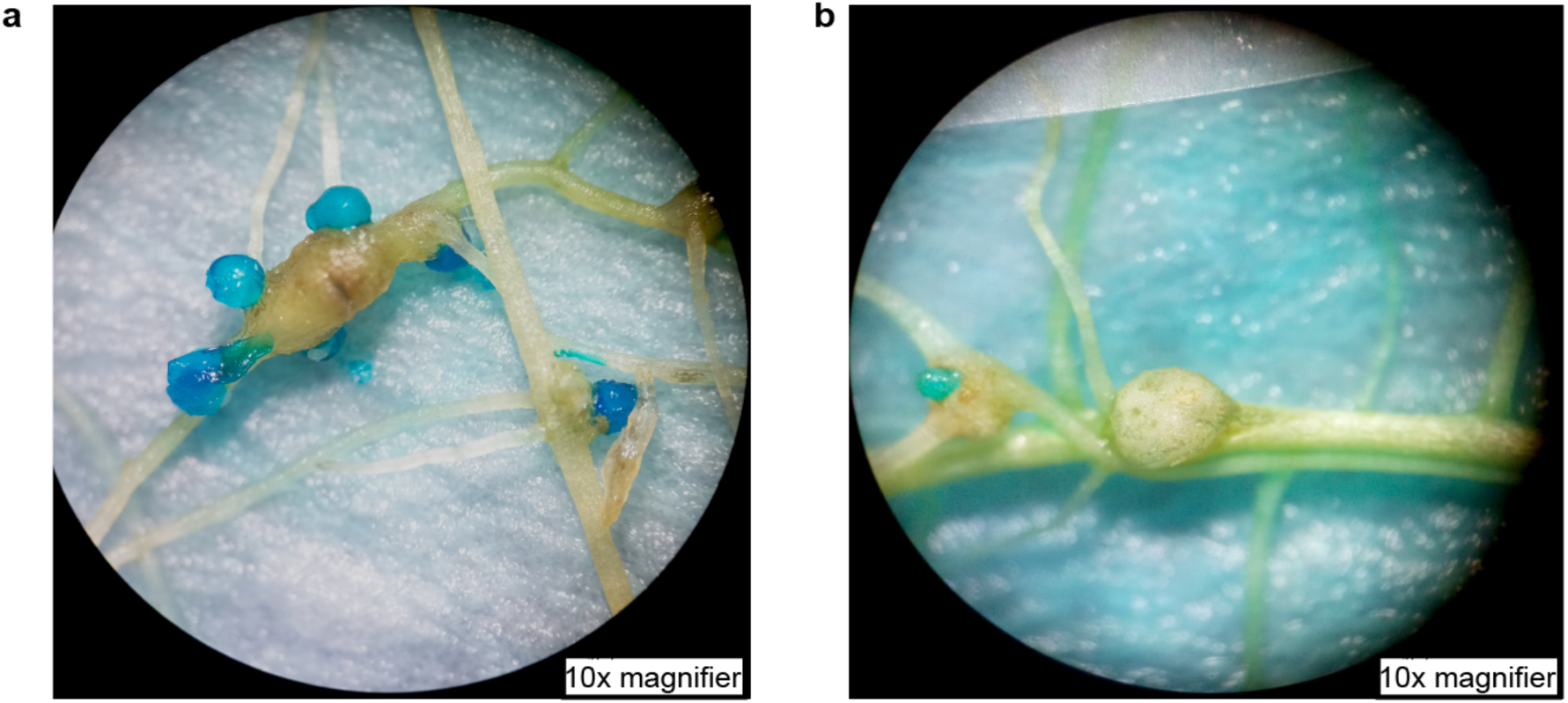
Phenotypic evaluation of root-knot nematode (RKN) infection in common bean. **(a)** Panel shows an egg mass (EM) stained with eriglaucine, highlighting RKN oviposition on the root surface. **(b)** Shows a root gall, an indicator of nematode-induced tissue hypertrophy. Both traits, egg mass (EM) and root-gallong index (RI), were assessed as part of the phenotyping to evaluate resistance levels among genotypes. Images captured under a 10x magnifier.

